# Plug-and-Play Metabolic Transducers Expand the Chemical Detection Space of Cell-Free Biosensors

**DOI:** 10.1101/397315

**Authors:** Peter L Voyvodic, Amir Pandi, Mathilde Koch, Jean-Loup Faulon, Jerome Bonnet

## Abstract

Cell-free transcription-translation systems have great potential for biosensing, yet the range of detectable chemicals is limited. Here we provide a framework to expand the range of molecules detectable by cell-free biosensors by combining synthetic metabolic cascades with transcription factor-based networks. These hybrid cell-free biosensors are highly-sensitive and have a fast response and high-dynamic range. This work provides a foundation to engineer modular cell-free biosensors tailored for many applications.

## MAIN TEXT

There is currently an urgent need for low-cost biosensors in a variety of fields from environmental remediation to clinical diagnostics^1–3^. The ability of living organisms to detect signals in their environment and transduce them into a response can be utilized to create cheap, novel sensors with high sensitivity and specificity. By leveraging the ability of transcription factors to control gene expression, synthetic biologists have genetically engineered microbes to detect a wide range of compounds, from clinical biomarkers to environmental pollutants^4–7^.

Cell-free transcription/translation (TXTL) systems have great promise as the next generation of synthetic biology-derived biosensors. They are cheap to produce^8^, abiotic, and can be lyophilized such that they are stable at room temperature for up to one year: a vital necessity for point-of-care applications such as low-resource nation and home diagnostic use^9^. Cell-free TXTL toolboxes have been designed that support the operation of many of the circuits previously engineered *in vivo*^10,11^ and existing cell-free biosensors can successfully detect Zika virus in rhesus macaques and an acyl homoserine lactone, 3OC12-HSL, from *Pseudomonas aeruginosa* in human clinical samples^12,13^. However, current cell-free biosensors have been limited to detection of nucleic acid sequences, via toehold displacement, or well-characterized transcription factor ligands. Here we put forward a generalized, modular framework utilizing metabolic transducers to rapidly expand the chemical space detectable by cell-free biosensors in a plug-and-play manner.

Synthetic metabolic cascades have been used by the synthetic biology community for a wide range of applications, including production of biofuels, pharmaceuticals, and biomaterials^14–16^. As such, there is a wide variety of well-characterized enzymes catalyzing various reactions transforming one molecule into another. Our framework harnesses this power by using metabolic enzymes as transducers to allow us to ‘plug in’ a given enzyme into our characterized biosensor modules to detect a ligand with no known transcription factor analog (**Fig. 1a**). Specifically, the metabolic enzyme converts the undetectable molecule into one for which we have an existing transcription factor-based genetic circuit (**Fig. 1b**). We used the SensiPath webserver that we previously designed and validated *in vivo* to determine the required metabolic cascade^17,18^.

**Figure 1:**
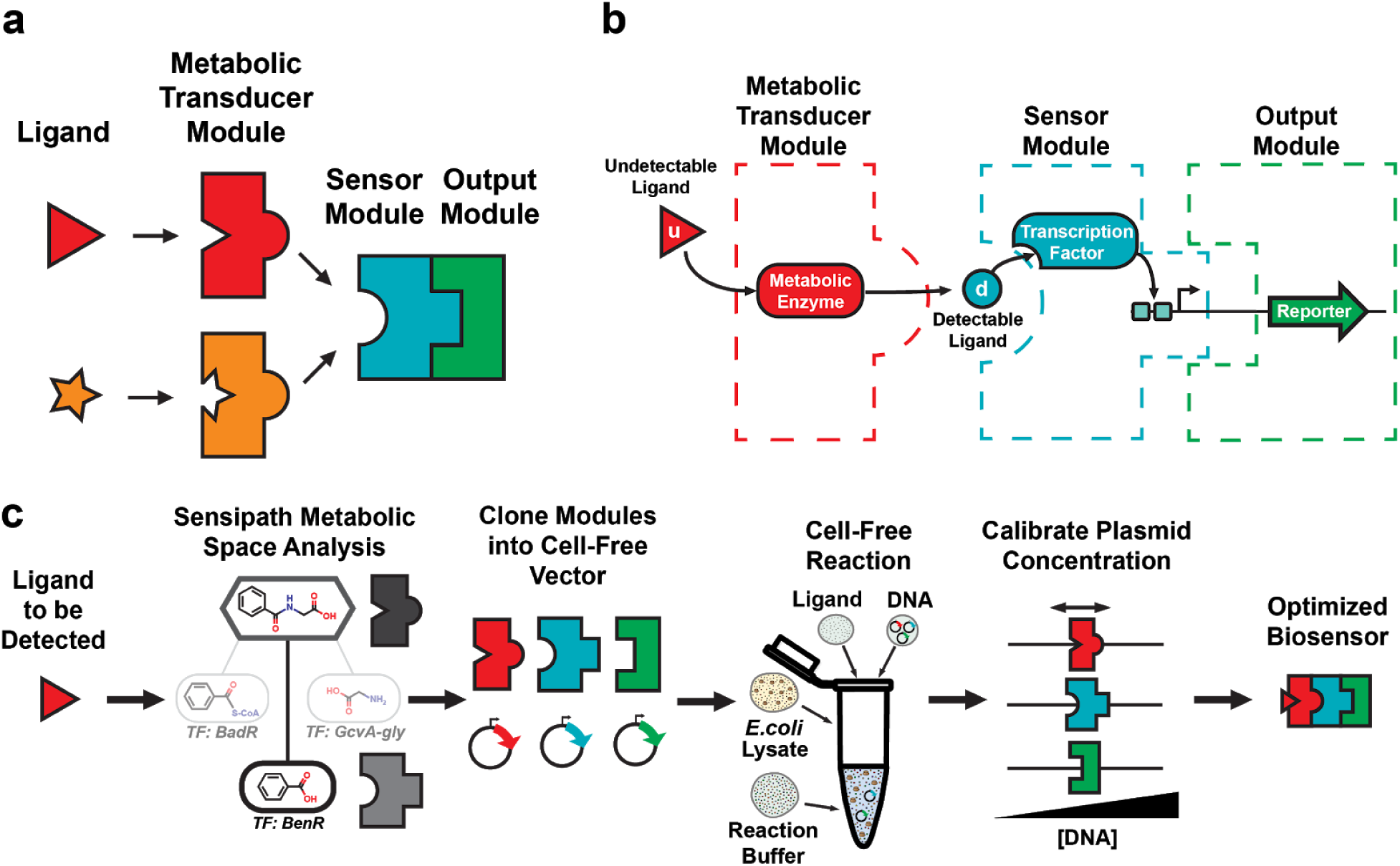
A modular design framework for engineering scalable cell-free biosensors. (**a**) Cell-free biosensors are composed of three modules: a generic sensor module linked to an output module and a metabolic transducer module transforming different molecules into ligands detectable by the sensor module. (**b**) An undetectable ligand is converted into a detectable ligand by the enzyme from the transducer module. Binding to the transcription factor controls the sensor module and downstream gene expression. (**c**) The biosensor design workflow starts with retrosynthetic pathway design using the SensiPath server^18^. Once the transducer and sensor modules are determined, the genes encoding enzymes, transcription factors, and target promoters driving a reporter are cloned into cell-free expression vectors. The sensor is calibrated by titrating the concentrations of each plasmid to maximize signal output and dynamic range.

The workflow to engineer a cell-free biosensor detecting a novel molecule is straightforward **(Fig. 1c)**. First, possible metabolic pathways to convert the molecule of interest into a detectable ligand are identified using SensiPath. Second, the genes coding for the metabolic transducer enzyme, the transcription factor (TF) sensor, and the reporter module are synthesized and cloned into cell-free expression vectors. Finally, the DNA concentration of each plasmid is titrated in cell-free reactions to optimize signal strength and dynamic range in response to the molecule of interest (**Fig. 1c**).

As a proof-of-concept example of this system, we engineered a sensor for benzoic acid using the transcription factor BenR and expanded its detection capabilities with two different metabolic transducers: one for hippuric acid using the HipO hippurase and one for cocaine using the CocE esterase.

BenR is a member of the AraC/XylS family of transcription factors, originally from *Pseudomonas putida*. In the presence of benzoate, BenR binds to the Ben promoter and activates transcription (**Fig. 2a**). To engineer a benzoate cell-free biosensor, we cloned BenR under the control of the OR2-OR1-Pr promoter, a modified version of the lambda phage repressor promoter Cro, known to express strongly in cell-free systems^19^. The P_Ben_ promoter driving super-folder green fluorescent protein (sfGFP) was cloned in a separate plasmid. After initial pilot tests demonstrated that BenR was functional in a cell-free environment, we optimized the BenR biosensor by titrating the DNA concentration of the TF and reporter plasmids.

**Figure 2:**
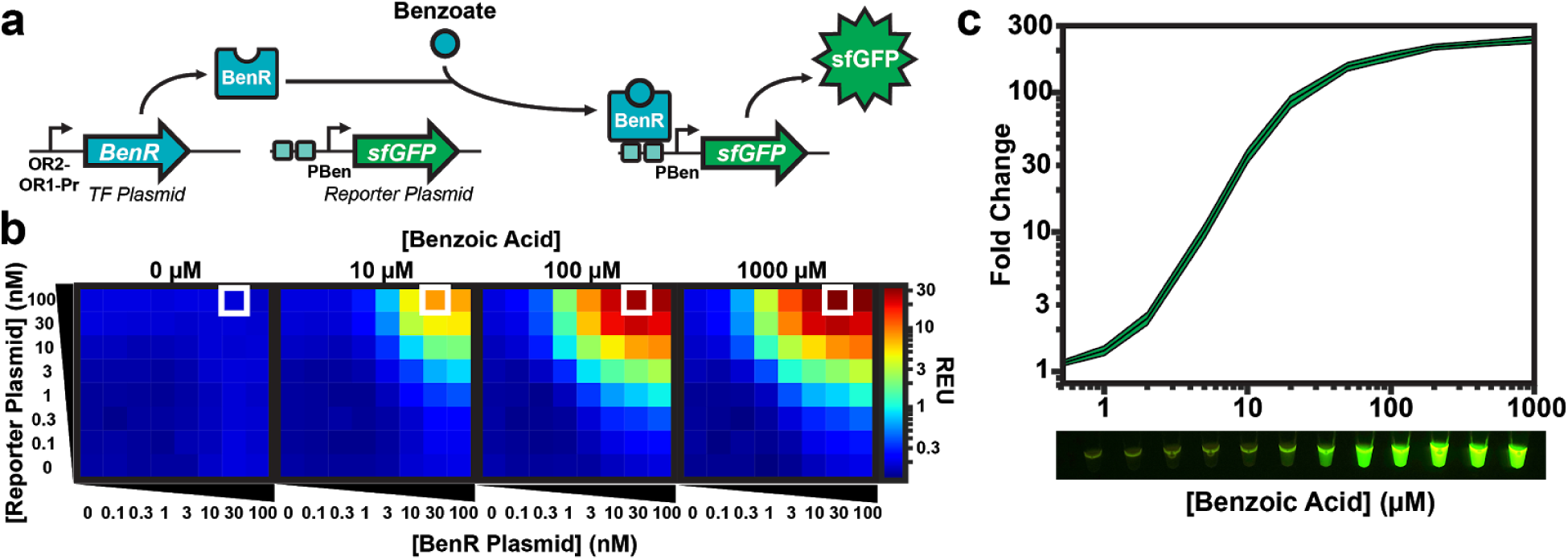
Calibration of sensor and output modules for benzoate detection. (**a**) BenR binds to the PBen promoter in the presence of benzoate and activates gene expression. Here BenR is cloned in the pBEAST plasmid (a derivative of pBEST^19^) and driven by a strong constitutive promoter, OR2-OR1-Pr. The P_Ben_ promoter is cloned into another pBEAST backbone and drives expression of the superfolder green fluorescent protein (sfGFP). Because the system operates without a cellular boundary, multiple plasmids encoding different components of the network can easily be used simultaneously. Plasmids concentrations can then be fine tuned to identify optimal operating conditions. (**b**) Optimization of the BenR sensor and reporter modules. Cell-free reactions of 20 µl containing different concentrations of the BenR and reporter plasmids were prepared and their response to different concentrations of benzoic acid were monitored. The white square represents the optimal condition (100 nM reporter and 30 nM BenR plasmid) with the highest-fold change and higher sensitivity. (see **Supplementary Fig. 1 and Supplementary Table 1**). Reactions were run in sealed 384 well-plates in a plate-reader at 37°C for at least eight hours. The heat maps represent the signal intensity after four hours. Data are the mean of three experiments performed on three different days and all fluorescence values are expressed in Relative Expression Units (REU) compared to 100 pM of a strong, constitutive sfGFP-producing plasmid. See methods for more details. (**c**) *Upper panel*: The BenR sensor can detect benzoic acid over three orders of magnitude and at concentrations as low as 1 µM. Shaded area around curves corresponds to +-SD from the mean of the 3 experiments. *Lower panel*: GFP expression in response to the same range of concentrations of benzoic acid as in the upper panel is easily detectable by eye on a UV table.

One advantage of working in a cell-free framework is that the DNA concentration is directly controlled by pipetting. As such, the process of finding an optimal DNA concentration is relatively straightforward: we created a matrix of DNA concentrations for TF and reporter plasmids between 0 nM and 100 nM and induced these different cell-free reactions using four different concentrations of benzoic acid: 0 µM, 10 µM, 100 µM, and 1000 µM (**Fig. 2b, Supplementary Table 1**).

Encouragingly, the system had extremely low background signal in the absence of benzoic acid, indicating that the P_Ben_ promoter has very little ‘leakiness’ in a cell-free environment. When benzoic acid was added to the reaction, the sfGFP output signal was clearly detectable and fluorescence intensity was correlated with increasing reporter plasmid concentration. However, the signal reached a plateau for increasing concentrations of TF plasmid at 30 nM. We hypothesize that this plateau is due to competition for transcriptional and translational resources between transcription factor and reporter plasmid. This plateau is also observed in a mathematical model of cell-free biosensors (**Supplementary text**). Based on this data, we set the optimal plasmids concentrations to 30 nM for the TF plasmid and 100 nM for the reporter plasmid.

Compared to its *in vivo* counterpart^17^, the cell-free benzoic acid biosensor is faster (maximum signal reached in four hours, **Supplementary Fig. 1**) and far more sensitive (**Fig. 2c**). In addition, the cell-free biosensor version has a much higher dynamic range and a maximum fold change of over 200 (vs. ∼10-fold *in vivo*). These results exemplify the advantages of cell-free systems for rapidly engineering biosensors with optimal properties.

With the sensor and output modules optimized, we demonstrated the ability of our system to expand its chemical detection space using different metabolic transducer modules. HipO is an enzyme from *Campylobacter jejuni* that converts hippuric acid into benzoic acid while CocE is an esterase from *Rhodococcus sp.* that converts cocaine into benzoic acid. We cloned each enzyme into the cell-free expression vector and, using the optimized DNA concentrations of TF and reporter plasmids, titrated different concentrations of metabolic transducer DNA for a range of inducer inputs (**Fig. 3a, Supplementary Table 2**). Interestingly, we observed a clear peak in sfGFP signal corresponding to a particular concentration effectiveness: 3 nM for HipO and 10 nM for CocE. We built several mathematical models based on different assumptions that could reproduce the observed bell-shaped response to enzyme DNA concentration as well as its shift between the two enzymes (**Supplementary Fig. 2**). Based on these models, we concluded that the observed bell-shaped response is most likely due to competition between the different modules, leading to an important and unnecessary enzyme production at high DNA concentrations that divert resources such as RNA polymerase, ribosomes, and energy from sfGFP transcription and translation, as well as generating toxic byproducts. Moreover, we identified that the shifting peak between the two setups is most probably due to lower expression of CocE (**Supplementary text and Supplementary Fig. 3**).

**Figure 3:**
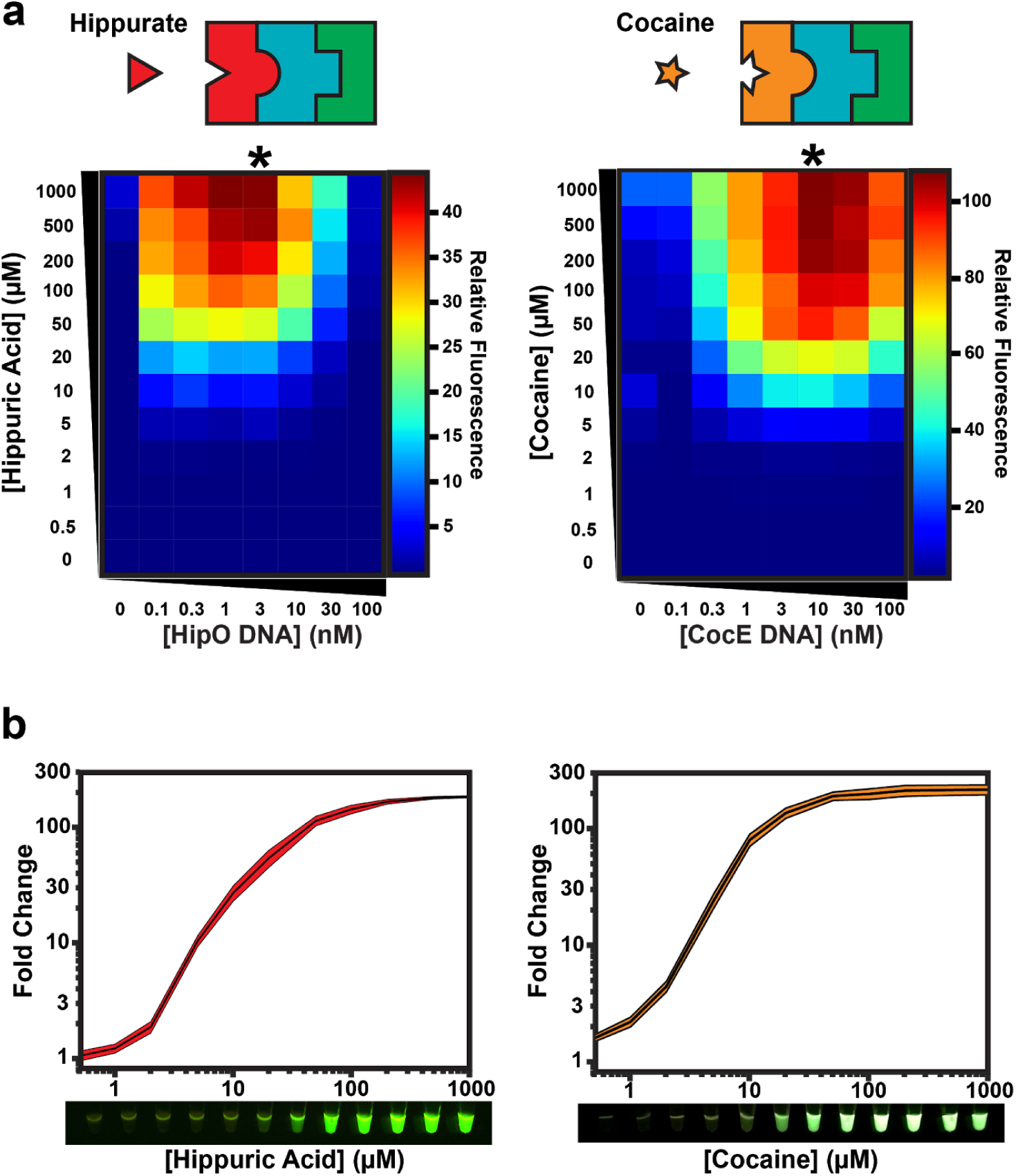
Expanding the chemical detection space of cell-free biosensors by plugging various metabolic transducers into an optimized sensor module. (**a**) Hippurate or cocaine can be detected using different metabolic transducers. Plasmids encoding the HipO or CocE enzymes, which convert hippuric acid or cocaine into benzoic acid, were mixed at different concentrations with optimal BenR and reporter plasmids concentrations as determined in **Fig. 2** (30 nM and 100 nM, respectively). These reactions were then incubated with increasing concentrations of inducer for at least eight hours. The heat maps represent the signal intensity after four hours (**Supplementary Fig. 4-5 and Supplementary Table 2**). Asterisks denote the optimal DNA concentration for the metabolic module. Data are the average of three experiments performed on three different days and all fluorescence values are expressed in Relative Expression Units (REU) compared to 100 pM of a strong, constitutive sfGFP-producing plasmid. (**b**) Optimized cell-free biosensors incorporating a metabolic transducer module exhibit comparable performance to the BenR sensor module (from **Fig. 2c**). All data are the mean of three experiments performed on three different days. Shaded area around curves corresponds to +-SD from the mean of the 3 experiments. See methods for more details. *Lower panel*: GFP expression in cell-free reactions in response to various concentrations of inducer visualized on a UV table.

A key observation is that even at very high levels of inducer, there is very little signal in the absence of DNA encoding the metabolic transducer. These data indicate that the metabolic enzyme is essential for sensor selectivity and that hippuric acid and cocaine have minimal off-target binding to BenR. Strikingly, both the hippuric acid and cocaine biosensors exhibit fold change and detection range highly similar to that of the benzoic acid sensor, demonstrating the high conversion rate of the metabolic transducer. The conversion also appears to be extremely fast as no significant difference was observed in responses kinetics with or without the metabolic transducer (**Supplementary Fig. 1, 4 and 5)**.

This work demonstrates that we can engineer modular, cell-free biosensors that can be easily calibrated to have high signal strength and dynamic range. Upon engineering a novel cell-free biosensor for benzoic acid, we show that the system can be scaled by using different metabolic transducer modules to expand the chemical space that each sensor/reporter pair can detect. In addition, we provide a rigorous scheme to optimize cell-free biosensor performance along with a mathematical model enabling a better understanding of the parameters governing cell-free biosensors response which will help future optimisation of such devices. Using our design framework, this process should be applicable to a wide range of other sensor/reporter pairs. We computed that 1205 disease associated biomarkers from the Human Metabolome Database (HMDB) could be converted into a detectable molecules by one enzymatic reaction, of which 64 biomarkers that could be transformed into benzoate and thus theoretically connected via a metabolic transducer to our optimized sensor (see **Supplementary Text**). By rapidly expanding the number of detectable compounds, cell-free biosensors using plug-and-play metabolic transducers can address many challenges such as environmental detection, drug enforcement, and point-of-care medical diagnostics.

## Acknowledgments

We thank the Bonnet and Faulon lab members for fruitful discussion. We are grateful to Pau Bernado, Annika Urbanek, and Anna Morato for helpful discussions on cell-free systems. This work was supported by an ERC starting grant “COMPUCELL” to JB. JB also acknowledges continuous support from the INSERM Atip-Avenir program and the Bettencourt-Schueller Foundation. The CBS acknowledges support from the French Infrastructure for Integrated Structural Biology (FRISBI) ANR-10-INSB-05-01. JLF acknowledges support from BBSRC/EPSRC (grant number BB/M017702/1) and from the ANR (grant number ANR-15-CE1-0008). MK is supported by DGA (French Ministry of Defense) and Ecole Polytechnique and AP is supported by a scholarship from the IDEX Paris-Saclay. All plasmids will be available from Addgene.

## Author contributions

P.L.V., A.P., J-L.F. and J.B. designed experiments, P.L.V. and A.P. cloned constructs and performed experiments, and M.K. constructed the computer model simulations. All of the authors contributed to the writing of the manuscript.

## Data availability statement

Source data for main text figures, along with details on models, and DNA sequences for all constructs are provided in supplementary materials. All other raw data are available from the corresponding authors on reasonable request.

## SUPPLEMENTARY MATERIALS

This article contains 5 supplementary files.

1. **Supplementary file 1.** **Supplementary Information:** supplementary text, figures and tables: **- Supplementary Text:** **-** SensiPath Metabolic Space Analysis **-** Mathematical Model of Cell-Free Biosensors **- Supplementary Fig. 1**: Time course of the benzoic acid biosensor response to varying concentrations of inducer. **- Supplementary Fig. 2**: Modeling metabolic transducer behavior for HipO and CocE **- Supplementary Fig. 3**: Superfolder-GFP expression with J23101 and pBEST promoter (OR2-OR1-Pr). **- Supplementary Fig. 4**: Time course of the hippuric acid biosensor response to varying concentrations of inducer. **- Supplementary Fig. 5**: Time course of the cocaine biosensor response to varying concentrations of inducer. **- Supplementary Table 1:** Fluorescence results from calibration of TF and reporter plasmids. - **Supplementary Table 2:** Fluorescence results from calibration of HipO and CocE metabolic transducer plasmids
2. **2. Supplementary file 2.** Mathematical-model derivation.
3. **Supplementary file 3.** DNA sequences: **-** DNA sequences.zip containing: **-** pBEAST-BenR.gb **-** pBEAST-PBen-sfGFP.gb pBEST-HipO.gb **-** pBEAST-J23101-CocE.gb **-** pBEAST-sfGFP.gb **-** pBEAST-J23101-sfGFP.gb
4. **Supplementary file 4.** <u>Supplementary table 3</u>: List of biomarkers extracted from the HMDB database, the effectors that can directly detect those biomarkers, and the biomarkers that can be transformed via a metabolic reaction into a detectable molecules, along with the associated metabolic reaction and corresponding enzymes.
5. **Supplementary file 5.** <u>Supplementary table 4</u>: List of biomarkers that can be transformed into benzoate through a metabolic transducer, along with the metabolic reactions and the associated enzymes.

## MATERIALS AND METHODS

### Molecular biology

All clones were based on a previously characterized cell-free expression plasmid (pBEST-OR2-OR1-Pr-UTR1-deGFP-T500 was a gift from Vincent Noireaux [Addgene plasmid # 40019]^19^). To better facilitate cloning with a range of techniques and any future component insertion into larger gene circuits, the construct was modified by adding 40 base pair spacers and an upstream terminator and renamed pBEAST. Clones were created via Gibson or Golden Gate assembly in DH5αZ1 chemically competent *E. coli* where the deGFP was replaced by BenR or HipO. For CocE, the promoter was changed to another strong constitutive promoter, J23101, and RBS, B0032. The reporter plasmid for P_Ben_ used native RBS from *Pseudomonas putida* and superfolder-GFP as the output, which was found to give a stronger, faster signal in cell-free reactions at 37°C. DNA for cell-free reactions was prepared from overnight bacterial cultures using Maxiprep kits (Macherey-Nagel). Plasmids used in this paper will be available from Addgene.

### Extract preparation

Cell-free *E. coli* extract was produced using a modified version of existing protocols^8,20^. An overnight culture of BL21 Star (DE3) *E. coli* was used to inoculate 660 mL of 2xYT-P media in each of six 2 L flasks at a dilution of 1:100. The cultures were grown at 37°C with 220 rpm shaking for approximately 3.5 hours until the OD 600 = 2.0. Cultures were spun down at 5000 x g at 4°C for 12 minutes. Cell pellets were washed twice with 200 mL S30A buffer (14 mM Mg-glutamate, 60 mM K-glutamate, 50 mM Tris, pH 7.7), centrifuging afterwards at 5000 x g at 4°C for 12 minutes. Cell pellets were then resuspended in 40 mL S30A buffer and transferred to pre-weighed 50 mL Falcon conical tubes where they were centrifuged twice at 2000 x g at 4°C for 8 and 2 minutes, respectively, removing the supernatant after each. Finally, the tubes were reweighed and flash frozen in liquid nitrogen before storing at −80°C.

Cell pellets were thawed on ice and resuspended in 1 mL S30A buffer per gram cell pellet. Cell suspensions were lysed via a single pass through a French press homogenizer (Avestin; Emulsiflex-C3) at 15000-20000 psi and then centrifuged at 12000 x g at 4°C for 30 minutes to separate out cellular cytoplasm. After centrifugation, the supernatant was collected and incubated at 37°C with 220 rpm shaking for 60 minutes to digest remaining mRNA with endogenous nucleases^8^. Subsequently, the extract was recentrifuged at 12000 x g at 4°C for 30 minutes, and the supernatant was transferred to 12-14 kDa MWCO dialysis tubing (Spectrum Labs; Spectra/Por4) and dialyzed against 2 L of S30B buffer (14 mM Mg-glutamate, 60 mM K-glutamate, ∼5 mM Tris, pH 8.2) overnight at 4°C. The following day, the extract was re-centrifuged at 12000 x g at 4°C for 30 minutes. The supernatant was optionally concentrated using a 10,000 MWCO centrifuge column (GE Healthcare; Vivaspin20) based on total protein levels from a Bradford assay (ThermoScientific) to obtain concentrations above 15 mg/mL, aliquoted, and flash frozen in liquid nitrogen before storage at −80°C.

### Cell-Free Reactions

Cell-free reactions were prepared by mixing 33.3% cell extract, 41.7% buffer, and 25% plasmid DNA, any inducer, and water. Buffer composition was made such that final reaction concentrations were as follows: 1.5 mM each amino acid except leucine, 1.25 mM leucine, 50 mM HEPES, 1.5 mM ATP and GTP, 0.9 mM CTP and UTP, 0.2 mg/mL tRNA, 0.26 mM CoA, 0.33 mM NAD, 0.75 mM cAMP, 0.068 mM folinic acid, 1 mM spermidine, 30 mM 3-PGA, and 2% PEG-8000. Additionally, the Mg-glutamate (0-6 mM), K-glutamate (20-140 mM), and DTT (0-3 mM) levels were serially calibrated for each batch of cell-extract for maximum signal. Benzoic acid, hippuric acid, and cocaine hydrochloride were purchased from Sigma-Aldrich. Permission to purchase cocaine hydrochloride was given by the French drug regulatory agency (Agence Nationale de Sécurité du Médicament et des Produits de Santé) to allow development of a new biosensor. Inducers were dissolved in ethanol and final reactions contained 0.5% ethanol for all inducer concentrations including the negative control. Reactions were prepared in PCR tubes on ice and 20µL were transferred to a black, clear-bottom 384 well plate (ThermoScientific), sealed, and the reaction was carried out in a plate reader (Biotek; Cytation3 or Synergy HTX) to measure both endpoints and reaction kinetics. The subsequent data were processed and graphs created using custom Python scripts or Microsoft Excel. Reactions for the representative images in **Figure 2c** and **Figure 3b** were incubated in PCR tubes at 37°C for four hours and imaged on a UV table with either a Sony *α*6000 camera (benzoic and hippuric acid sensors) or a cell phone camera (cocaine sensor) and background subtracted with Adobe Photoshop.

### Reaction Models

Coarse-grained modeling was performed using ordinary differential equations, simulated using the R software. Briefly, the model combines Michaelis-Menten kinetics for the adaptor module and resource competition for RNA polymerases and ribosomes to account for varying DNA concentration effects. Hill equations are used for promoter activation. Production of toxic byproducts as well as energy consumption for protein production were also included. Full model derivation can be found in the supplementary materials.

### Code availability

Simulation scripts are available at https://github.com/brsynth. Custom python scripts used to process data are available upon request to the authors.

